# Bi-directional electron transfer between H_2_ and NADPH mitigates the response to light fluctuations in green algae

**DOI:** 10.1101/2020.10.08.331504

**Authors:** Yuval Milrad, Shira Schweitzer, Yael Feldman, Iftach Yacoby

**Affiliations:** School of Plant Sciences and Food Security, The George S. Wise Faculty of Life Sciences, Tel Aviv University, Ramat Aviv, Tel Aviv 69978, Israel

**Author notes:** Corresponding author: Iftach Yacoby, Corresponding author. Contributed equally (Yuval Milrad, Shira Schweitzer). Author contributions: YM, SS and IY design and coordinated the study. YM, SS carried out the biochemical studies, photosynthetic parameters, performed the MIMS measurements and drafted the manuscript. YF participated in the photosynthetic parameter’s analysis. IY conceived the study, coordinated the research and drafted the manuscript. All authors read and approved the final manuscript. Funding: The research was funded by NSF-BSF energy for sustainability 2016666 and by ISF 1646/16.

**Keywords:** Photosynthesis, Hydrogen, Chlamydomonas, Anaerobic metabolism, NADPH

## Abstract

The metabolism of green algae has been the focus of much research over the last century. These photosynthetic organisms can thrive under various conditions and adapt quickly to changing environments by concomitant usage of several metabolic apparatuses. The main electron coordinator in their chloroplasts, nicotinamide adenine dinucleotide phosphate (NADPH), participates in many enzymatic activities and is also responsible for interorganelle communication. Under anaerobic conditions, green algae also accumulate molecular hydrogen (H_2_), a promising alternative for fossil fuels. However, in order to scale-up its accumulation, a firm understanding of its integration in the photosynthetic apparatus is still lacking. While it is generally accepted that NADPH metabolism correlates to H_2_ accumulation, the mechanism of this collaboration is still vague and rely on indirect measurements. Here, we investigated this connection using simultaneous measurements of both dissolved gases concentration, NADPH fluorescence and electrochromic shifts at 520-546 nm. Our results indicate that energy transfer between H_2_ and NADPH is bi-directional and crucial for the maintenance of redox balance under light fluctuations. At light onset, NADPH consumption is initially eventuated in H_2_ evolution, which initiate the photosynthetic electron flow. Later on, as illumination continues the majority of NADPH is recycled by Nda2 rather than consumed by terminal sinks such as CBB cycle and H_2_ production. Dark onset triggers re-assimilation of H_2_, which produces NADPH and so, enables initiation of dark fermentative metabolism.

**One sentence summary:** Energy transfer between H_2_ and NADPH is bi-directional and crucial for the maintenance of redox balance under light fluctuations.

## Introduction

Nicotinamide adenine dinucleotide phosphate (NADPH) is an important electron carrier in the chloroplasts of photosynthetic organisms. It participates in many metabolic pathways as an electron donor and mediates interactions with other organelles (Burlacot et al., 2019). NADPH is produced mainly by the enzyme ferredoxin-NADPH oxidoreductase (FNR), while FNR itself is reduced by the single electron carrier; ferredoxin (Hemschemeier and Happe, 2011). The reduction of NADP^+^ due to light absorbance by photosystem I (PSI) is considered to be the major sink of electrons that are generated by the photosynthetic apparatus. It was postulated that the binding of FNR to the acceptor site of PSI grants it this superiority (Yacoby et al., 2011; Marco et al., 2019). Although the pool of NADPH is mainly assimilated by the Calvin-Benson-Basham (CBB) cycle, it can also be oxidized by the Plastoquinone (PQ) pool. In green algae, a type II NADPH dehydrogenase (Nda2) was shown to mediate this interaction with the PQ pool (Desplats et al., 2009), thus contributing reducing power back to the photosynthetic chain in an ‘alternative cyclic electron flow’ (Mus et al., 2005). This form of electron flow generates a proton gradient, which promotes ATP synthesis. However, in contrast to linear electron flow (LEF) it does not alter the pool of the reduced form of NADPH.

In green algae, as anoxia eliminates respiration and triggers decreased levels of ATP, the balance between ATP and NADPH is shifted and the activity of CBB cycle is hindered (Allen, 2003). This imbalance also generates a state in which NADP^+^ reduction is not preferred and thus induces acceptor side limitation on PSI (Clowez et al., 2015). Accordingly, H_2_ production by hydrogenase at light onset alleviates these limitations and contributes to a build-up of proton gradient, which supports ATP production (Ghysels et al., 2013). As a result, following dark incubation, H_2_ accumulates rapidly at the onset of light. However, this accumulation lasts only for a short period of time, after which hydrogenase is inactivated due to a shift of the electron flow toward other metabolic pathways. This shift has recently been the topic of some studies (Cournac et al., 2002; Godaux et al., 2015; Burlacot et al., 2018; Milrad et al., 2018), and is considered as the major limitation for sustained photosynthetic H_2_ harvesting (Tóth and Yacoby, 2019). Indeed it was suggested that NADP^+^ reduction outcompetes H_2_ evolution, however these postulations were based on the measurements of other metabolites such as CO_2_ and O_2_. Therefore, to date, a conformation to the connection between the metabolisms of NADPH and H_2_ is still missing.

Another process which was postulated to connect H_2_ and NADPH metabolism is the re-assimilation of H_2_ by the algae (Maione and Gibbs, 1986; Chen and Gibbs, 1992). Accordingly, electrons from H_2_ are transferred through hydrogenase back to ferredoxin, which in turn reduces NADP^+^ via FNR. Such process is beneficial to the algae since the accumulated H_2_, which diffuses out of the cells, generates a loss of potential energy. This loss perplexed researchers over the years, as in early studies, it was demonstrated that CO_2_ reduction takes place under anaerobic conditions, while pure H_2_ atmosphere serves as an electron donor, defined as photoreduction (Gaffron 1939). Indeed, H_2_ uptake was observed in *C. reinhardtii* both in light (Kosourov et al., 2012; Noone et al., 2017; Milrad et al., 2018) and in the dark (Scoma and Hemschemeier, 2017). However to date, there is no concrete evidence which might correlate the kinetics of NADP^+^ reduction and H_2_ assimilation.

Here, we investigated the connection between reduction/oxidation kinetics of NADPH and H_2_. To do so, we, combined a membrane inlet mass spectrometer (MIMS) with a dual pulse amplitude modulated fluorometer (Dual-PAM) and monitored the total electron flow in the thylakoid membrane by measuring the electrochromic shifts (ECS), in a Joliot type spectrometer (JTS). We exposed *C. reinhardtii* cells to light following dark anaerobic incubation and examined the effect it had on the kinetics of metabolites such as H_2_, O_2_, CO_2_ and NADPH. Our results suggest that the relationship between H_2_ and NADPH is bi-directional and contributes to the ability of the cells to gain balance in the ratio of ATP:NADPH.

## Results

### Algae metabolism under fluctuating light

In this work, we studied the reduction/oxidation kinetics of various metabolites which take part in the photosynthetic apparatus as well as the electrochromatic shift at 520-546 nm (ECS). To do so, we combined gas exchange measurements using a membrane inlet mass spectrometer (MIMS) (Liran et al., 2016; Milrad et al., 2018; Ben-zvi et al., 2019) and a dual pulse amplitude modulated fluorometer (DUAL-PAM-100, Walz) equipped with a DUAL-NADPH/9-AA module (Schreiber and Klughammer, 2009) besides we also used a Joliot type (JTS-100) spectrophotometer. This setup enabled us to simultaneously monitor the concentrations of dissolved gasses such as H_2_, O_2_ and CO_2_, in parallel to PSII and NADPH fluorescence (Figure. 1). In addition, it granted us the ability to determine the rates of various electron flow pathways. All measurements were conducted under anoxia, which was achieved by dark incubation for an hour in a sealed 5 ml cuvette. In order to test the effect of light on these metabolites, we exposed the cells to 10 repeated cycles of illumination (370 µE m^−2^ s^−1^) and darkness of two and three minutes, respectively. We have previously shown that such a scheme did not alter the enzymatic pool of hydrogenase and that cycles 2 to 10 feature an identical phenotype (Milrad et al. 2018, see also Figure. S1). Therefore, in order to decrease background noises, we averaged the data of these cycles. In addition, to confirm that the effects on the NADPH signal originated from biological processes, we subtracted the signal of a cell-free medium from the obtained results (see Figure. S2). The results of this experiment on cells of the wild-type (w.t) strain CC400-cw15 are shown in (Figure. 1; dashed red lines). At the establishment of dark anoxia, the cells did not fixate CO_2_, and H_2_ assimilation was slow. Accordingly, NADPH signal was stable at a ‘high steady state’. At the onset of light, H_2_ was immediately produced at a rate of 22.4 ± 3.4 e^-^ PS^-1^ sec^-1^. Its accumulation rate then decreased as CO_2_ began to fixate in a net rate of 11.2 ± 0.4 e^-^ PS^-1^ sec^-1^, as we previously reported (Milrad et al., 2018). Concurrently, NADPH signal decreased exponentially with an initial rate of 27.7 ± 3.4 e^-^ PS^-1^ sec^-1^, until it reached a ‘low steady state’ in which the net NADPH was stable again. At dark onset, we observed an expected O_2_ respiration along with a decrease in the concentration of the accumulated H_2_ (at a rate of 12.4 ± 1.5 e^-^ PS^-1^ sec^-1^). Furthermore, CO_2_ levels increased for a minute, until its concentration was stabilized. Concomitantly, NADPH signal dropped sharply at the onset of darkness (at a maximal rate of 264 ± 4 e^-^ PS^-1^ sec^-1^). However, this decrease was very brief; within a few seconds the signal shifted, and NADPH began to accumulate at around 22 e^-^ PS^-1^ sec^-1^ in an increasing rate (up to 65 e^-^ PS^-1^ sec^-1^) for a minute. As it reached the concentration of the ‘high steady state’ the accumulation abruptly ceased, and the levels of NADPH remained stable.

**Figure.1:**
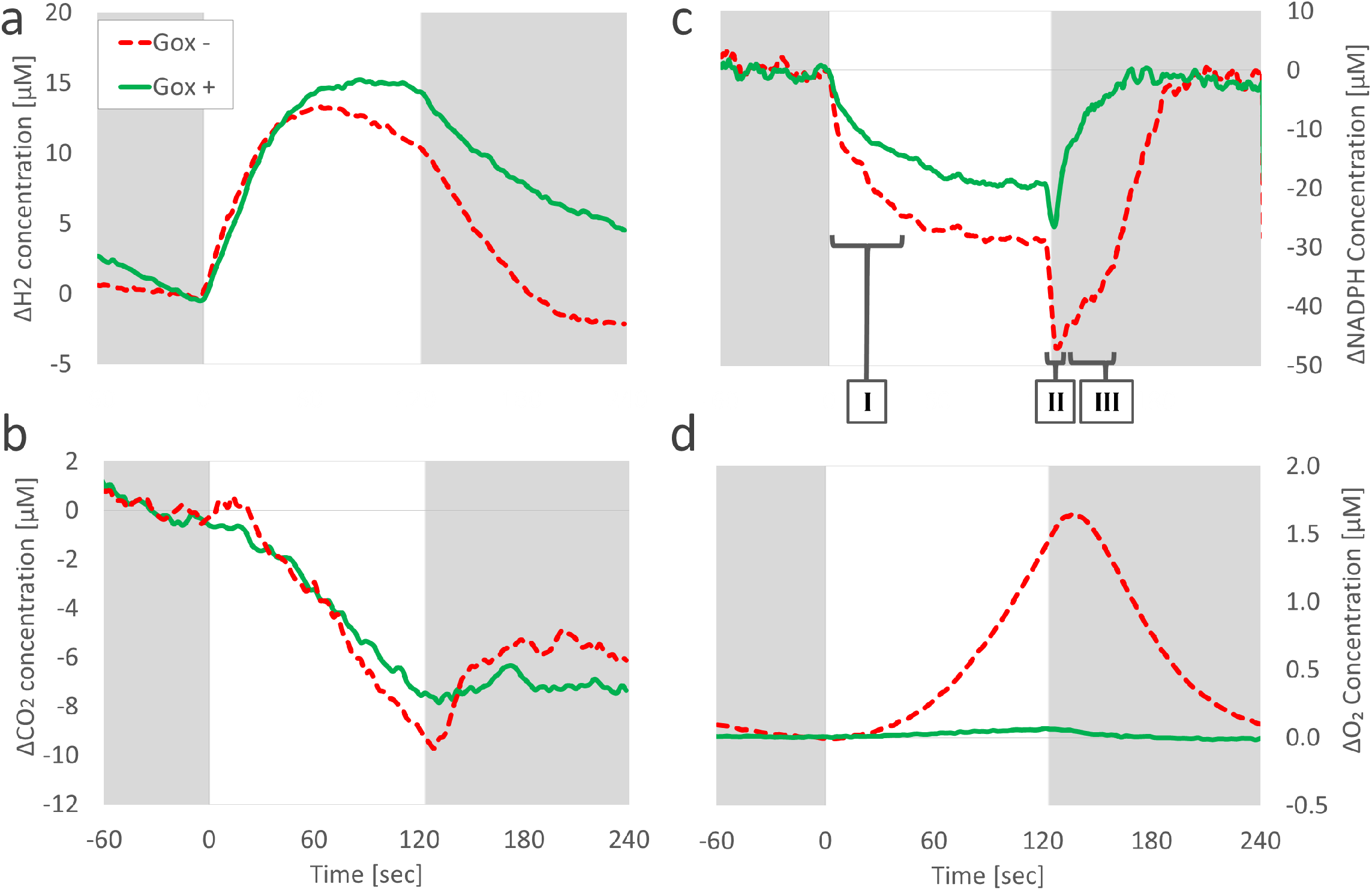
Light/dark fluctuations kinetics in the presence or absence of O_2_ Scavenger: *C. reinhardtii* wild-type strain (CC400-cw15) was incubated for 1 h in the dark in the absence (red; dashed line) or presence (green; solid line) of an oxygen scavenger system: glucose oxidase, glucose and catalase. After incubation, the cells were exposed to 10 cycles of 2 minutes of illumination (370 µE m^−2^ s^−1^; white background) and 3 minutes of darkness (gray background). H_2_ (a), CO_2_ (b), NADPH (c) and O_2_ (d) concentrations were measured using a membrane inlet spectrometer (MIMS) combined with a Dual-PAM. The values are the mean of cycles 2 to 10 of three biological replications. 3 domains were examined (underlined in c), for each the electron transfer rates were calculated using a linear regression: (I) Light onset, (II) NADPH drop at dark onset and (III) dark acclimation.

### Alternations in metabolism caused by strict anoxia

Since O_2_ can be utilized by many participants in the chloroplast such as the FLV proteins, as well as in the mitochondria, we could not isolate the pathway which causes the observed shifts in NADPH levels. Therefore, we neutralized the activity of FLV proteins as well as the Mehler reaction in PSI, by adding the O_2_ scavenger glucose oxidase (GOx), supplied with glucose and catalase (Roberty et al., 2014). We then repeated the light/dark cycle regime and compared the results (Figure. 1; solid green lines). The comparison between these treatments are presented along the text and are shown in (Table 1). We have previously shown that GOx can efficiently remove intracellular O_2_ (Milrad et al., 2018) and indeed no O_2_ accumulation was observed in the experiment. We observed that removing O_2_ hardly altered the H_2_ production (20.3 ± 2.9 e^-^ PS^-1^ sec^-1^) nor the fixation rates of CO_2_ (8.3 ± 1.7 e^-^ PS^-1^ sec^-1^), as we reported in (Milrad et al., 2018). It did, however, decrease the decay of NADPH signal at light onset (from 27.7 ± 3.4 e^-^ PS^-1^ sec^-1^ in presence of O_2_ to 19.4 ± 1.1 e^-^ PS^-1^ sec^-1^ in its absence). In addition, at the onset of darkness, H_2_ uptake rate was slower in the presence of GOx (from a rate of 12.4 ± 1.5 e^-^ PS^-1^ sec^-1^ in presence of O_2_ to rate of 8.9 ± 1.5 e^-^ PS^-1^ sec^-1^ in its absence) and the cessation in CO_2_ fixation did not feature any increase. Interestingly, the sharp drop in NADPH signal was halved (132 ± 12 e^-^ PS^-1^ sec^-1^) and its increase kinetics back to the ‘high steady state’ was (33.5 ± 2.1 e^-^ PS^-1^ sec^-1^). Still, the evolution stopped as it reached the ‘high steady state’, a minute following the onset of darkness.

**Table.1:**
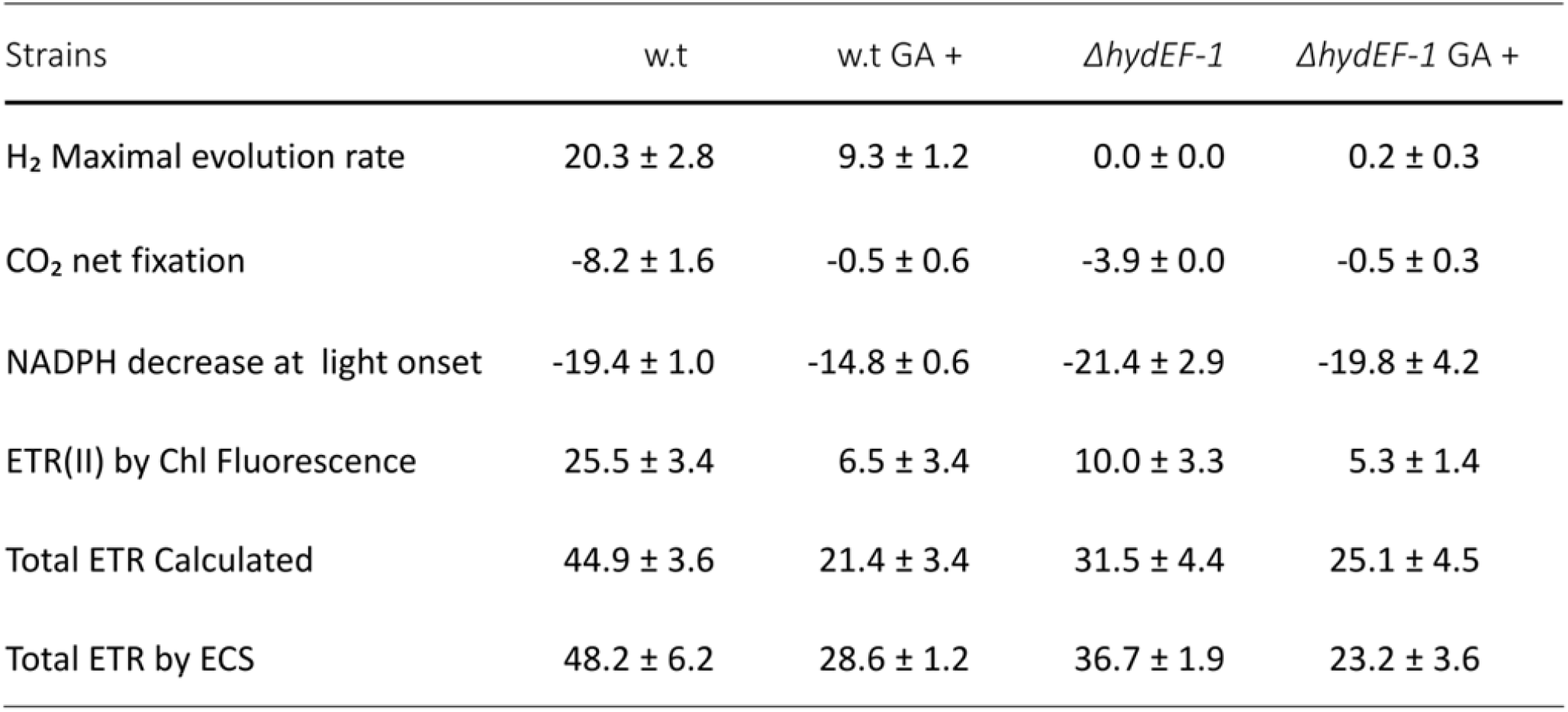
Calculated electron flow: The electron flux generated by light exposure (370 μE m^-2^ sec^-1^), as presented in Figure. 2, was calculated as described in material & methods for each of the major sinks (H_2_ evolution and CO_2_ fixation). In addition, the initial decrease of NADPH concentration, which can be attributed to the activity of Nda2, indicates a contribution of the alternative electron flow. PSII fluorescence was simultaneously measured and used to calculate the contribution of the linear electron flow by ETR(II). Summation of the linear and the alternative electron flow provide the total electron flux during light exposure. Interestingly, the calculated rate matches the total electron flow rate which was measured by the ECS-dirk protocol. The table present the results of either the wild-type (w.t) strain CC400-cw15 of Δ*hydEF-1*, to which glucose oxidase, catalase and glucose were added, in the absence or presence of glycaolaldehyde (GA). All of the results are presented in e PS^-1^ sec^-1^. The values are the mean of three biological replications, with calculated standard error.

### The effect of hydrogenase on NDPH kinetics

To test the role of hydrogenase activity in NADPH metabolism, we compared the w.t strain CC400-cw15 to *ΔhydEF-1*, a mutant lacking an active hydrogenase (Posewitz et al., 2004). Since it was reported that the absence of active hydrogenase affects the photosynthetic activity, and thus the levels of accumulated O_2_ (Ghysels et al., 2013), we conducted the comparison under strict anoxia by an addition of GOx. For the w.t strain, we use the results in the presence of GOx shown in (Figure. 1). The combined results are shown in (Figure. 2; solid lines). As expected, at light onset, H_2_ production (Figure. 2a) was not detected in *ΔhydEF-1*, and the rate of net CO_2_ fixation (Figure. 2b) was lower (4.0 ± 0.1 e^-^ PS^-1^ sec^-1^). The NADPH signal, in the absence of active hydrogenase, featured a rapid increase as a response to light onset (at a rate of 143 ± 16 e^-^ PS^-1^ sec^-1^), which was absent in the w.t strain. This initial increase in *ΔhydEF-1* instantly shifted direction to an exponential decrease (at a maximal rate of 21.5 ± 3.0 e^-^ PS^-1^ sec^-1^) until the ‘low steady state’ level was reached. Since no oxygen was accumulated in the experiment, we used PSII fluorescence (Figure. 2d, e) and electrochromatic shift (ECS) at 520-546 nm (Figure. 2f) as a monitoring system for the indication of linear electron flow from PSII. Our results featured the same phenotype for both strains: An increase of the PSII fluorescence at light onset, followed by a slow decrease as the cells adapt to the irradiance. In addition, when we exposed the cells to saturating pulses prior and during illumination (Figure. 2d), we observed that the potential activity of PSII is intact, as there are no differences in the dark adapted Fv/Fm (Figure. 2e). However, the electron transfer rate of PSII during illumination was diminished in *ΔhydEF-1* (From 25.5 ± 3.5 e^-^ PS^-1^ sec^-1^ to 10.1 ± 3.4 e^-^ PS^-1^ sec^-1^ Figure. 2 e, f). At dark onset (120 sec), the sharp drop in NADPH signal was considerably less prominent in *ΔhydEF-1* (89 ± 15 e^-^ PS^-1^ sec^-1^ versus 132 ± 12 e^-^ PS^-1^ sec^-1^ in the w.t, as was the increase back to the ‘high steady state’ (19.6 ± 4.9 e^-^ PS^-1^ sec^-1^). These observations suggest a linkage between hydrogen uptake and NADP^+^ reduction at dark onset. To verify whether H_2_ uptake results from a biological activity or a physical phenomenon, we injected H_2_ saturated standard into a cell-free medium and observed the changes in H_2_ concentration. We also repeated this assay with dark-adapted cultures of both a w.t strain (D66) and another strain lacking hydrogenase activity (*ΔhydA1-1 hydA2-1* double mutant) (Figure. S3). The results show that following a rapid stabilization (for ∼2 minutes, due to dissolution of the highly concentrated H_2_ standard injection), H_2_ consumption takes place only in the presence of active hydrogenase.

**Figure.2:**
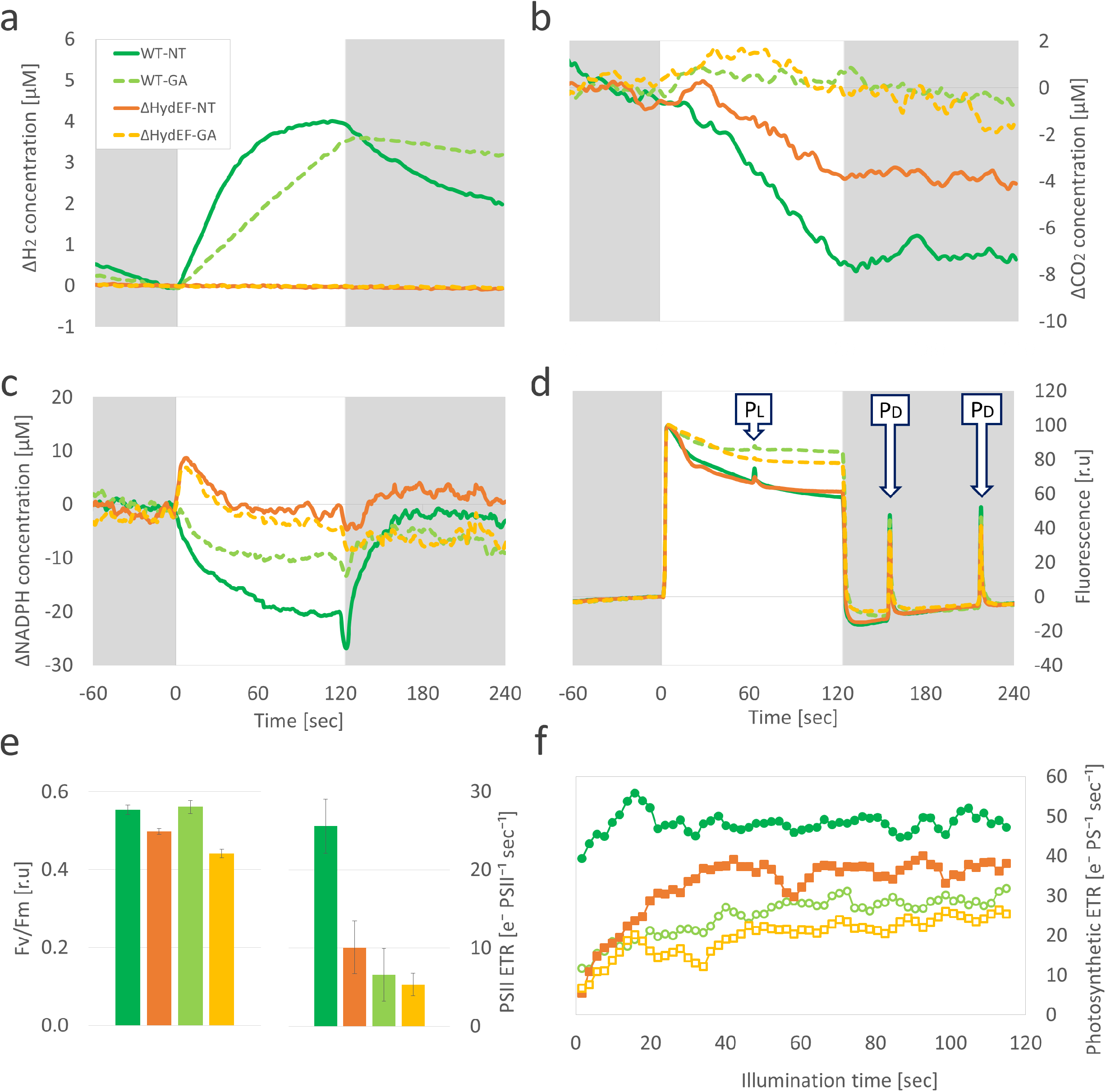
Light/dark fluctuations kinetics in the absence of Hydrogenase: *C. reinhardtii* wild-type (WT; green) or Hydrogenase maturation mutant (*ΔhydEF-1*; orange) were incubated for an hour in the dark, in the presence of an oxygen scavenger system: glucose oxidase, glucose and catalase. After incubation, the cells were exposed to 10 cycles of 2 minutes of illumination (370 µE m^−2^ s^−1^; white background) and 3 minutes of darkness (gray background). H_2_ (a) and CO_2_ (b) concentrations were measured using a membrane inlet spectrometer (MIMS), simultaneously to fluorescence measurements of NADPH (c) and photosystem II (d) using a Dual-PAM-100. During (P_L_) and between (P_D_) each illumination cycle, the cells were exposed to saturating light pulses (10,000 µE m^−2^ s^−1^ for 200 ms, see panel d). Fv/Fm values (left) and PSII electron transfer rate (right) were calculated (e). In addition, the total photosynthetic electron transfer rate during the illumination was monitored by implementing DIRK protocol (f). The cells were tested in the absence of inhibitors (NT; solid lines) or in the presence of glycolaldehyde (GA; dashed lines in a-d or open markers in f). The values are the mean of cycles 2 to 10 (in a-e) or cycles 2-4 (in f) of three biological replications.

### CBB inactivation in the presence/absence of active hydrogenase

Due to the fact that uptake of H_2_ coincides with NADPH increase, we examined the involvement of CBB cycle, a major sink for NADPH, in H_2_ uptake. To do so, we blocked CBB cycle by adding Glycolaldehyde (GA), a known inhibitor of phosphoribulokinase (Roberty et al., 2014) to both strains (*ΔhydEF-1* and CC400-cw15). Results from the light/dark cycle regime with GA are shown in (Figure. 2; dashed lines). In addition, the rates which were calculated for both strains, in the presence or absence of GA were summarized in (Table. 1). In the presence of GA and light, carbon fixation of the w.t strain was inhibited and its H_2_ accumulation was linear at a rate of 9.3 ± 1.2 e^-^ PS^-1^ sec^-1^, as we previously described (Milrad et al., 2018). In addition, in the w.t strain, the decrease of NADPH at light onset was slightly slower in the presence of GA (14.9 ± 0.7 e^-^ PS^-1^ sec^-1^) versus in its absence (19.4 ± 1.1 e^-^ PS^-1^ sec^-1^). Interestingly, the initial increase of NADPH at light onset in *ΔhydEF-1* was not altered in the presence of GA (150 ± 16 e^-^ PS^-1^ sec^-1^) *versus* in its absence (143 ± 16 e^-^ PS^-1^ sec^-1^). Furthermore, the kinetics of NADPH oxidation during light (decrease rate of 21.5 ± 3.0 e^-^ PS^-1^ sec^-1^) was also similar in the presence of GA (maximal rate of 19.9 ± 4.3 e^-^ PS^-1^ sec^-1^). These results imply that the unique rise of NADPH at light onset, seen only in *ΔhydEF-1*, is not related to the CBB cycle. It should be noted that samples treated with GA showed a smaller decrease in PSII fluorescence (Figure. 2d) and lower rate of PSII activity (Figure. 2e,f), regardless to the fact that its potential activity was not altered (as the Fv/Fm values were not changed, Figure. 2e). These observations can be inferred as lower level of photosynthetic activity due to limitations on PSI acceptor side. Alternatively, the total electron transfer was directly measured using the ECS-DIRK method at 520-546nm (Bailleul et al., 2010). Methodically, during illumination, we exposed the cells to short dark pulses (of 60 ms). At the onset of darkness, the absorbance drops immediately, and the decrease rate during 10ms, (following an initial 2 ms of the decrease) was fitted using a linear regression to give a rate constant. To calculate the actual electron flow, this constant was divided by the maximal absorbance at 520-546nm obtained following a single red saturating laser (Figure. 2f) (Bailleul et al. 2010, see also Figure. S5). The results indeed show that the rate of total electron transfer directly measured by ECS-DIRK method (48.3 ± 6.3 e^-^ PS^-1^ sec^-1^) matches our combined chlorophyll and NADPH fluorescence decrease at light onset estimations (45.0 ± 3.6 e^-^ PS^-1^ sec^-1^). The same can be observed for the total electron transfer rate in the presence of GA. While the combined chlorophyll and NADPH fluorescence decrease at light onset estimations resulted in an electron flow rate of 21.5 ± 3.5 e^-^ PS^-1^ sec^-1^, the rate which was gained using the electrochromatic shift measurements was 28.7 ± 1.3 e^-^ PS^-1^ sec^-1^. In addition, the electron transfer rate that was calculated for the *ΔhydEF-1* mutant by combining the rates of NADPH fluorescence decrease at light onset and PSII chlorophyll fluorescence were 31.5 ± 4.5 and 25.2 ± 4.5 e^-^ PS^-1^ sec^-1^ in the absence/ presence of GA respectively and the rates that were measured by the electrochromatic shifts were 36.8 ± 2.0 and 23.2 ± 3.6 e^-^ PS^-1^ sec^-1^ in the absence/ presence of GA respectively. Another interesting phenomenon was observed at the onset of darkness; the presence of GA diminished the sharp decrease of NADPH to 41 ± 16 e^-^ PS^-1^ sec^-1^ in the w.t (compared to 132 ± 12 e^-^ PS^-1^ sec^-1^ for the untreated cells), but not in *ΔhydEF-1*, which had a rate of 97 ± 13 e^-^ PS^-1^ sec^-1^ (compared to 89 ± 15 e^-^ PS^-1^ sec^-1^ for the untreated cells). Furthermore, during darkness and in the presence of GA, NADPH levels in the ‘high steady state’ were also lower in both strains. Interestingly, H_2_ consumption rate at dark onset was much lower in the presence of GA (Figure. 2a). These results coincide with the observations of the NADPH signal, implying that the differences within NADPH oxidation/ reduction kinetics are likely due to hydrogenase activity.

### The imbalance of ATP:NADPH ratio hinders NADPH metabolism

To gain further insights into the role of ATP:NADPH balance on the kinetics of H_2_ and NADPH shifts, we tested the effect of known photosynthetic inhibitors (Figure. 3). For a clear comparison, we also present the traces of the untreated w.t strain, as were shown in (Figure. 1; solid green lines). The addition of the PSII acceptor side inhibitor 3-(3,4-dichlorophenyl)-1,1-dimethylurea (DCMU) gravely decreased H_2_ evolution. In fact, H_2_ evolution lasted only few seconds (at a rate of 5.2 ± 0.9 e^-^ PS^-1^ sec^-1^) before it was completely hindered, a phenomena which was also reported in previous works (Cournac et al., 2002). It should be noted that the CBB cycle is hindered but not inactivated under such conditions. Interestingly, the NADPH signal did decrease exponentially (at a rate of 13.3 ± 0.4 e^-^ PS^-1^ sec^-1^), in a manner which resembled the effect of GA addition (Figure. 2c). Moreover, as the light was turned off, a sharp decrease of NADPH signal (at a rate of 63 ± 14 e^-^ PS^-1^ sec^-1^) followed by a rise back to the ‘High steady state’ were observed, also as in the presence of GA. These observations may imply that the changes in NADPH levels are not solely dependent on carbon fixation, but rather by an imbalanced redox potential promoted when carbon fixation or PSII activity are damaged. In order to elucidate the influence of redox potential on NADPH kinetics, we studied the effects of uncoupler carbonyl-cyanide *m*-chlorophenyl-hydrazine (CCCP) addition. Under these conditions, we observed a sharp drop in NADPH levels at light onset, yet no further decrease was detected and the ‘low steady state’ was achieved immediately. In addition, the sharp decrease at the dark onset was eliminated, and the reduction of NADPH back to the ‘high steady state’ was immediate. Concurrently, H_2_ evolved in a linear manner under light (at a rate of 8.7 ± 0.9 e^-^ PS^-1^ sec^-1^), and no H_2_ consumption was detected when the light was turned off.

**Figure.3:**
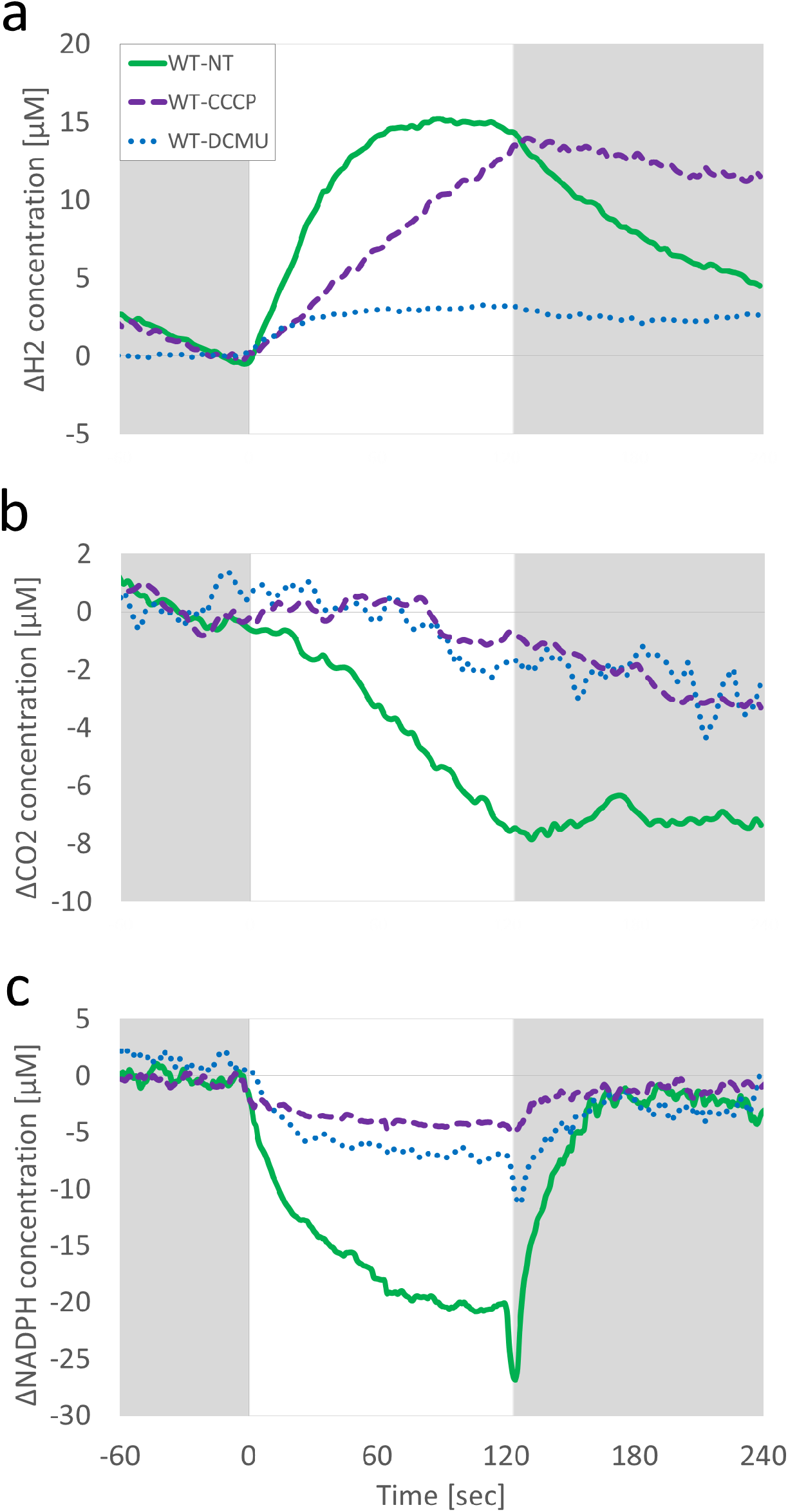
Light/dark fluctuations kinetics in the presence of photosynthetic inhibitors: *C. reinhardtii* wild-type were incubated for 1 hour in the dark in the presence of an oxygen scavenger system: glucose oxidase, glucose and catalase. After incubation, the cells were exposed to cycles of 2 minutes of illumination (370 µE m^−2^ s^−1^; white background) and 3 minutes of darkness (gray background). H_2_ (a) CO_2_ (b) and NADPH (c) concentrations were measured using a membrane inlet spectrometer (MIMS) combined with a Dual-PAM in the absence of inhibitors (solid green lines) or in the presence of either CCCP (dashed purple lines) or DCMU (dotted blue lines). The values are the mean of cycles 2 to 10 of three biological replications.

### The contribution of linear electron flow to terminal sinks

Following the data presented above, we wondered if there is an agreement between the rate of linear electron flow (LEF) and the reduction rates of the major terminal sinks; CBB cycle and H_2_ production. To test this, LEF, was measured independently via two different methods; either by conventional chlorophyll fluorescence or by applying the ECS-DIRK method, following a subtraction of the CEF contribution (see methods). As can be seen in (Figure. 4; yellow triangles vs blue circles) the two measurements are fairly in line with each other for both strains, *i*.*e*. the w.t (Figure. 4a) and the *ΔhydEF-1* (Figure. 4b) mutant. Next, we plotted the contribution of both sinks, CBB cycle and H_2_ production (Figure. 4 red and green respectively). Markedly a large variance was observed between the actual capacity of LEF at steady state (∼50 and ∼35 e^-^ PS^-1^ sec^-1^ for the w.t or *ΔhydEF-1*; respectively) *versus* the actual electron divergence for CBB cycle and H_2_ production combined (∼10 and ∼5 e^-^ PS^-1^ sec^-1^ for the w.t or *ΔhydEF*-1; respectively).

**Figure. 4:**
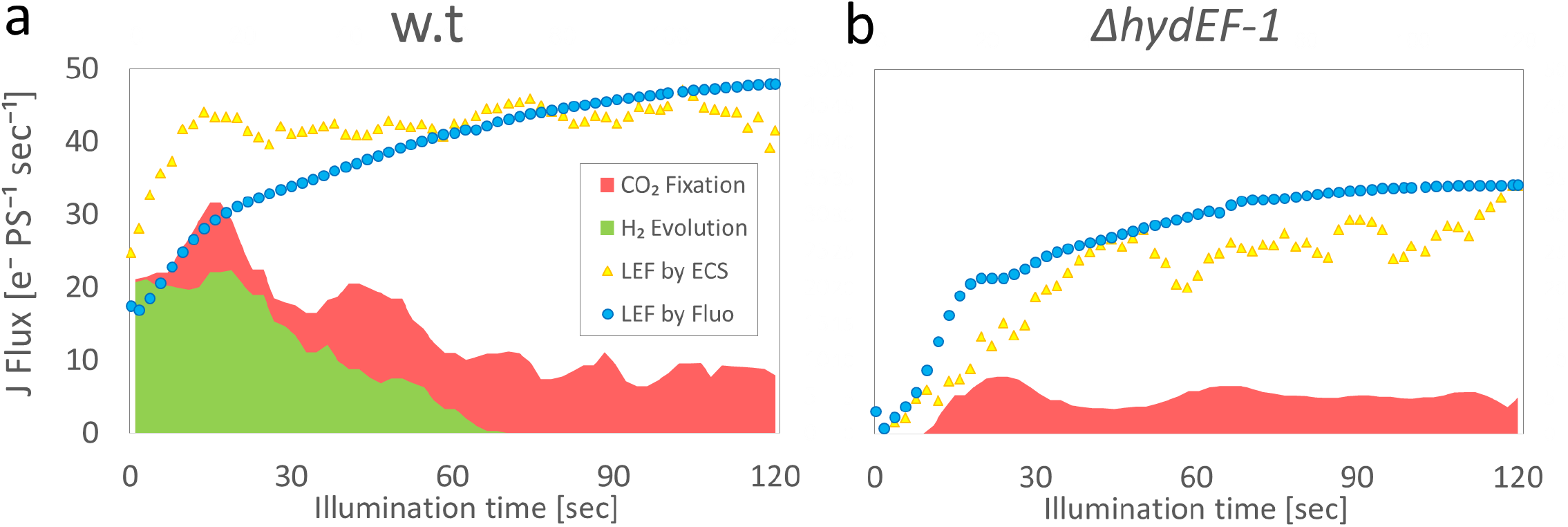
Linear electron flow versus the sinks: *C. reinhardtii* wild-type (w.t; a) or Hydrogenase maturation mutant (*ΔhydEF-1*; b) were incubated for an hour in the dark, in the presence of an oxygen scavenger system: glucose oxidase, glucose and catalase. After incubation, the cells were exposed to 10 cycles of 2 minutes of illumination and 3 minutes of darkness. The electrons consumed for H_2_ (green) production and CO_2_ fixation (red) were measured using a membrane inlet spectrometer (MIMS). Linear electron flow (LEF) was measured either by chlorophyll fluorescence using a Dual-PAM-100 (blue circles) or by DIRK-ECS protocol using JTS-100 spectrophotometer (yellow triangles). The values are the mean of cycles 2 to 10 (PAM) or cycles 2-4 (ECS-DIRK) of three biological replications.

## Discussion

In this work we simultaneously examined the kinetics of total electron flow, NADPH, H_2_ and CO_2_ metabolism, thus shedding light on the electron transfer between them on-line, in a w.t and in the *ΔhydEF-1* mutant. The resulting electron diversion map is shown in (Figure. 4). For the best of our knowledge, this is the first time in which such a map is presented. As will be discussed in detail below, we observed that the buildup of the ATP:NADPH ratio by hydrogenase is heavily dependent on the oxidation of the NADPH pool in Nda2. In turn, the balanced ATP:NADPH ratio enables NADP^+^ reduction by the photosynthetic apparatus under fluctuating light.

### Imbalanced ATP:NADPH ratio enhances NADPH consumption at light onset

Following dark incubation, the ratio between NADPH and ATP is imbalanced due to low ATP levels (Allen, 2003). This imbalance hinders the activity of CBB cycle, the main sink for NADPH, and it was previously shown to be activated only after a minute of illumination (Burlacot et al., 2018; Milrad et al., 2018). During this time, the imbalance is alleviated by processes which oxidize NADPH and concomitantly support ATP synthesis (Alric et al., 2010). Accordingly, we observed that NADPH concentration drops rapidly at the onset of illumination (Figure. 1c). Under aerobic conditions, this drop can be explained by O_2_ reduction taking place in Flavodiiron (FLV) proteins via NADPH oxidation (Chaux et al., 2017; Jokel et al., 2019). Indeed, an increase in NADPH consumption rate in the presence of accumulated O_2_ was observed, emphasizing the role of the FLV in alleviating the ATP:NADPH imbalance. However, FLVs are unlikely to account for the decrease in NADPH at light onset under strict anoxia. Since FLVs and the CBB cycle are yet to be activated, NADPH oxidation is likely performed by Nda2. Nda2 concomitantly reduces the PQ pool, which under anoxia can then reduce only Cytochrome_b6f_ (Cyt_b6f_), as chlororespiration requires O_2_ to function. Such scenario ultimately leads to generation of ATP via the resulting proton gradient and balances the ATP:NADPH ratio as the NADPH pool shrinks and the ATP pool enlarges.

### Hydrogenase activity alleviates the imbalanced ATP:NADPH ratio

NADPH oxidation in Nda2 also results in surplus electrons reaching PSI (Saroussi et al., 2016). These electrons are then distributed by ferredoxin to downstream sinks such as H^+^, which is reduced to molecular H_2_ by the enzyme hydrogenase (Winkler et al., 2009). Indeed, in the absence of an active hydrogenase, NADPH uptake at light onset did not take place (Figure. 2c; *ΔhydEF-1*). In fact, in *ΔhydEF-1* the opposite took place, and a fast rise in NADPH concentration was observed. These results are in agreement with previous observations, where H_2_ production rate at light onset was significantly impaired when Nda2 was inhibited (Mus et al., 2005). The combined electron transfer rate gained from measuring Nda2 activity and PSII ETR, lead to a similar number of total photosynthetic electron flow which was gained by direct measurement via the ECS-DIRK method, ∼45 e^-^ PS^-1^ sec^-1^ at light onset. Hence, at light onset H_2_ production consumes ∼50% of the total electron flow. Interestingly, the same fraction of electrons, ∼50% is allocated for H_2_ production in the presence of GA, which has a total electron pool of ∼25 e^-^ PS^-1^ sec^-1^. Hence, we suggest that at light onset about 50% of the electrons are in fact allocated to cyclic electron flow via the NADPH oxidation route. Furthermore, the addition of PSII inhibitor (DCMU) gravely limited electron flow but H_2_ evolution was nonetheless observed concomitantly to NADPH oxidation (Figure. 3). Interestingly, these two processes ceased at the same time, several seconds following light onset. These findings are in line with an electron flow from NADPH oxidation through Nda2 to H_2_ production at light onset.

### NADPH metabolism in the transition to dark anoxia

At dark onset, NADPH levels drop sharply as PSI activity halts and the reduction of NADP^+^ by FNR ceases. We suggest that the rapid drop in NADPH concentration at dark onset can report on the combined oxidation of NADPH by the CBB cycle, Nda2 and FLVs, all of which are not expected to immediately cease at dark onset. Indeed, the drop of NADPH signal at dark onset was smaller in the absence of O_2_, although CO_2_ fixation rates in the preceding light exposure were similar (Figure. 1b, c). Remarkably, an even further decrease in NADPH oxidation (smaller decrease in NADPH signal) was observed once the CBB cycle was inactivated by GA (Figure. 2). In the absence of hydrogenase, NADPH assimilation at dark onset was halved, probably due to minimal activation of the CBB cycle, as the ATP:NADPH ratio was imbalanced. Accordingly, an addition of CCCP destabilizes this ratio at low level, hinders the metabolism of NADPH, and eliminates the fast NADPH oxidation at dark onset. These results imply that the majority of the reduced NADPH is rapidly recycled back to NADP^+^, likely via Nda2 activity.

Following the fast NADPH oxidation at dark onset, both the consumption of NADPH and CO_2_ fixation cease (Figure. 1). Since it is unlikely that the entire pool of ATP, which was accumulated during the light, will be consumed immediately, one cannot help but wonder why the CBB cycle stops. Here, we show that when the CBB cycle was inactivated due to either direct inhibition or a disruption of the ATP:NADPH ratio, the transition from light to dark had a lesser effect on NADPH levels. Nevertheless, NADPH levels increased back to the ‘high steady state’, but never exceed it (Figure. 1). This observation can be explained by the need of the cells to maintain their reducing power. In contrast to light conditions, where photosynthetic organisms derive their energy and accumulate reducing power, to thrive under darkness they must reserve a reductant pool to allow sustainable dark fermentation metabolism (Catalanotti et al., 2013). Since in the absence of an active hydrogenase the restoration of NADPH pool at dark onset was the lowest (Figure. 2c), we suggest that H_2_ may serve as the major electron donor via hydrogenase to NADP^+^.

### H_2_ assimilation regulates the ATP:NADPH ratio in anoxia

It was previously suggested that H_2_ uptake hinders the potential of sustainable H_2_ production (Scoma and Hemschemeier, 2017; Kosourov and Jokel, 2018; Milrad et al., 2018). Such assimilation could be the result of a chemical oxy-hydrogen reaction, and indeed we observed elevated consumption of H_2_ in the presence of O_2_ (Figure. 1). However, the process of H_2_ uptake takes place only when hydrogenase is present (Figure. S3). In addition, we observed that once the CBB cycle was inhibited, H_2_ uptake was eliminated. However, we are unable to differentiate between the role of O_2_ reduction and CBB activity, since we conducted the experiments under strict anoxia. To get better insight on the subject, we reexamined the data from our previous work (Milrad et al., 2018). In that work, we tested the effects of CBB cycle inactivation in the absence of O_2_ scavengers, and indeed O_2_ accumulated at the same levels in all samples. Under these conditions, H_2_ uptake did not take place in the presence of O_2_. Therefore, we can state that the assimilation of H_2_ is dependent on CBB cycle activity. Furthermore, neither H_2_ uptake nor NADP^+^ reduction took place once the ATP:NADPH ratio was imbalanced (in the presence of the uncoupler CCCP, Figure. 3). Considering that H_2_ uptake and restoration of the NADPH pool coincided, it is likely that H_2_ serves as the electron donor, thus filling this void and balancing the reducing power for anoxic metabolism. The electron transfer from H_2_ toward NADP^+^ may imply an important role of H_2_ in balancing the NADP^+^/NADPH ratio.

## Conclusions

In this work we provided high resolution map of the electron divergence among multiple routes at light fluctuations. The rate of electron transfer for each of the processes as well as the total electron flow are presented in (Figure. 4). The white area encages between the total electron flow and the terminal sinks, the extremely high rate of NADPH consumption at dark onset and the extreme H2 production at light onset which is heavily dependent on NADPH oxidation by Nda2, suggest that NADPH recycling by Nda2 is in fact the major sink for NADPH oxidation.

## Materials & Methods

### Cell growth and conditions

*Chlamydomonas reinhardtii* wild type strains (CC400-cw15 and D66) or hydrogenase deficient mutants (*ΔhydEF-1* and *ΔhydA1-1 ΔhydA2-1*) were cultivated in 50 mL TAP medium (Tris/Acetate/Phosphate, see Appendix), kept in Erlenmeyer flasks capped with a silicone sponge. Cells were grown under constant irradiation of 60 µE m^−2^ s^−1^ at 24.5°C and stirring until they reached early log phase (2–5 µg Chl mL^−1^). The chlorophyll concentration was measured according to (Jeffrey and Humphrey, 1975).

### Experimental protocol

Early log phase cells were centrifuged at 4000 g for 2 min and resuspended to a final concentration of 15 µg Chl mL^−1^, in a medium containing TAP, 50 mM HEPES and 2 mM Na_2_CO_3_ at pH 7.2. The cells were then transferred to a sealed quartz cuvette covered with an aluminum foil to maintain darkness for 50 minutes. If stated, glucose oxidase (200 U ml^-1^), catalase (200 U ml^-1^) and glucose (50mM) (Roberty et al., 2014) were added at the beginning of the dark incubation, in order to eliminate oxygen evolution due to light exposure. Following incubation, the measurement cuvette was inserted into the Dual-PAM-100 cuvette holder (optical unit ED-101US/MD; Walz), which kept the sample thermostatic at 24.5°C during the experiment. At this point, if stated, either 60 mM Glycolaldehyde (GA) (Anderson et al., 2007), 40 μM 3-(3,4-dichlorophenyl)-1,1-dimethylurea (DCMU) (Roberty et al., 2014), or 20 μM carbonyl cyanide m-chlorophenylhydrazone (CCCP) (Cournac et al., 2002) were added. The cells were then kept under darkness for additional 10 minutes while connected to the measurement set-up, prior to the onset of illumination (370 µE m^−2^ s^−1^). Light intensity was determined by a Walz light detector (model US-SQS/L) attached to a Li-250A light meter (LI-COR Biosciences). Light and dark cycles (2 and 3 minutes respectively) were repeated 10 times and the data for cycles 2-10 was averaged. Due to analysis considerations, the data is presented in concentration changes. Therefore the results were shifted so time 0 represents the onset of light exposure and all changes are relative to the concentration which was determined at that point.

### Metabolites determination

Gas concentration within a liquid culture was determined as previously described in (Liran et al., 2016), using a membrane inlet mass spectrometer (MIMS). The measured masses were; 2, 32, 40 and 44 m/z, which correlate to the concentration of the molecules H_2_, O_2_, Ar and CO_2_, respectively. These different masses were detected using a 0.5 second dwelling time per mass. The traces of H_2_, O_2_ and CO_2_ were normalized using conversion factors, which were calculated from a standard curve of known gases concentration as described in (Liran et al., 2016). NADPH and chlorophyll fluorescence were obtained using a dual pulse amplitude modulated fluorimeter (DUAL-PAM-100; Heinz Walz Gmbh, Effeltrich, Germany). This fluorimeter possesses the properties of a pulse amplitude modulated (PAM) chlorophyll fluorimeter, with the addition of a dual wavelength absorbance spectrometer – the accessory NADPH/9-AA emitter-detector module (DUAL-D/ENADPH) (Schreiber and Klughammer, 2009). Using this module, NADPH fluorescence is excited at 365 nm using a UV-A Power LED and a photomultiplier sensor is employed for detection of blue-green emission (420 to 580 nm). The module is optimized for measuring light-induced changes of NADPH fluorescence in suspensions of isolated chloroplasts or algae, as previously described in (White et al., 2013). In order to eliminate scattering, we repeated the protocol for a cell-free TAP medium and used the data as a baseline (see Figure. S2). NADPH signal was converted to concentration by measuring the medium in darkness and injecting known amounts of NADPH. The signal was plotted and linearly fitted (see Figure. S4). Simultaneously, chlorophyll fluorescence was monitored using the DUAL-DPD detector. The measurement resolution was set at 10 ms and every 100 detections were averaged to obtain resolution of one second.

### Electron transfer rate determination

To calculate the rates of the increase/decrease of the metabolites, we fitted the curves in a linear regression. Then we divided the rate by the Chlorophyll concentration (15 μgr/ml in all experiments), divided by 2, assuming that half the chlorophylls are attributed to PSII ((Takahashi et al., 2013)) and then by the molaric mass of chlorophyll (894 gr/mol). We multiplied the result for the calculations of the electron flow toward H_2_ production, NADP^+^ reduction and NADPH oxidation by 2 or by 4 for the calculations of CO_2_ fixation rate. Finally, assuming that each PSI occupy 270 chlorophylls ((Takahashi et al., 2013)), we present the results in e^-^ PS^-1^ sec^-1^, as summarized in the following equation:

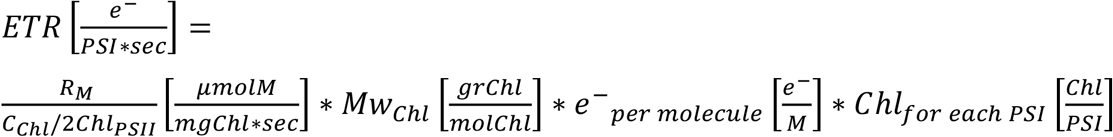

Where ***R***_***M***_ is the rate of the metabolite reduction/oxidation, ***C***_***Chl***_**/2*Chl***_***PSII***_ is the concentration of Chlorophyll in the sample divided by 2, ***Mw***_***Chl***_ is the molar mass of Chlorophyll, ***e***^−^_***per molecule***_ is the number of electron needed to reduce the metabolite and ***Chl***_***for each PSI***_ is the number of Chlorophylls in each PSI complex.

To assess PSII potential quantum yield (Y(II)) and ETR(II), saturating pulses (10,000 µE m^−2^ s^−1^ for 200ms) were fired during the dark periods (P_D_) and during illumination (P_L_). We observed that the Fm values were not altered by the light exposure, meaning that neither state transition nor non-photochremical quenching have a marked impact on ETR(II). We therefore, calculate the quantum yield as (Fm-Fs)/Fm for the whole duration of light exposure, as described in (Endo et al., 1995). The general photosynthetic electron transfer rate (ETR) was acquired by measuring the cells under the variant treatments, with the addition of 10% Ficoll to prevent cell sedimentation, in a Joliot-Type Spectrophotometer (JTS-100, Biologic). The JTS was supplied with a bi-lamp which measure the absorbance in either 520 or 546 nm, which enables fast and parallel measurement for the assessment of the electrochromatic shift (ECS), which was calculated by their subtraction (520-546 nm). Prior to illumination, we exposed the cells to a laser flash, provided by a dye laser (DCM, exciton laser dye) pumped by a frequency doubled Nd:YAG laser (Litron nano) and the photosynthetic charge recombination value was calculated from the initial shift as described in (Buchert et al., 2020). During the illumination, dark interval relaxation kinetics (DIRK) protocol (Bailleul et al., 2010) was implemented by exposing the cells to short dark pulses (60 ms) every 2 sec (total 60 pulses for each measurement, see Figure. S5). The rate of the decrease in the signal, as a result of this dark exposure, was subtracted from the shift of the steady state trace and divided by the value of the charge recombination.

## Acknowledgements

We would like to thank Eyal Dafni for Critical reading of this manuscript.

